# Early proteomic signatures of Alzheimer’s disease in the retina and brain of 3xTg-AD mice

**DOI:** 10.64898/2026.02.26.708380

**Authors:** Artjola Puja, Rachel McNeel, Rong Xu, Siyan Zhu, David Hansman, Jianhai Du

## Abstract

Visual dysfunction and retinal structural alterations often precede brain pathology and cognitive decline in Alzheimer’s disease (AD), yet the molecular basis of these early changes and their relationship to the brain pathology remain unclear. Here, we performed quantitative proteomic profiling of retina and brain from 1-month of age triple-transgenic (3xTg-AD) mice harboring human PS1^M146V^, APP^Swe^, and tau^P301L^ mutations, preceding detectable morphological abnormalities. Proteomic analysis identified 92 significantly altered proteins in the retina and 130 in the brain, with eight overlapping proteins between tissues. These shared proteins included three hemoglobin subunits (HBB1, HBB2, A8DUK4) and five proteins involved in metabolic regulation and intracellular transport. In addition to individual protein changes, pathway analysis demonstrated that mitochondrial metabolism and intracellular transport were commonly dysregulated in both tissues. Brain proteome was characterized by broad changes in mitochondrial-associated proteins, including respiratory chain components and mitochondrial ribosomal subunits, as well as proteins related to autophagy and synaptic vesicle pathways. In contrast, the retinal proteome was characterized by downregulation of vision-related proteins, altered small molecule transporters, and a marked reduction of the mitochondrial enzyme succinate-CoA ligase subunit β (SUCB2). As SUCB2 links mitochondrial metabolism to epigenetic regulation through succinylation and lactylation, its depletion may promote mitochondria-to-nucleus signaling and early transcriptional reprogramming in the AD retina. Together, these findings demonstrate early metabolic and transport dysregulation in both retina and brain and highlight selective alterations of visual proteins in the retina. These early retinal proteomic changes provide valuable insight into understanding early metabolic disturbances in the eye and brain for AD detection.

## Introduction

Alzheimer’s disease (AD) is a progressive neurodegenerative disorder and the leading cause of dementia worldwide (Perl, 2010; Rajan *et al*., 2021). Currently, there is no effective treatment to halt or reverse disease progression, and interventions started after clinical onset may contribute to the high failure of clinical trials (Dubois *et al*., 2014; Lane, Hardy and Schott, 2018; Kim *et al*., 2022). These limitations underscore the need to identify early biomarkers and define disease mechanisms during the preclinical stage of AD. The classical neuropathological hallmarks of AD are extracellular amyloid-β (Aβ) plaques and intracellular neurofibrillary tangles composed of hyperphosphorylated tau protein (Ratan *et al*., 2023). Aβ peptides, predominantly Aβ40 and the more aggregation-prone Aβ42, are generated from amyloid precursor protein (APP) through sequential proteolytic cleavage by β- and γ-secretases. Mutations in presenilin-1 (PS1), a core component of the γ-secretase complex, alter APP processing and preferentially increase Aβ42 production, thereby accelerating Aβ accumulation, synaptic dysfunction, and cognitive decline (Suzuki *et al*., 1994; Selkoe, 2000; Annaert and De Strooper, 2002; Sun *et al*., 2017). However, AD is now recognized as a multifactorial disorder that extends beyond Aβ and tau pathology. Emerging evidence highlights early mitochondrial and metabolic dysfunction, impaired proteostasis, neuroinflammation, and vascular contributions as integral components of disease progression (Di Filippo *et al*., 2010; Gadhave *et al*., 2021; Tian, Ji and Liu, 2022; Ottoy *et al*., 2023; Kazemeini *et al*., 2024). Importantly, AD-associated pathology is not restricted to the brain but also affects other regions of the central nervous system (CNS), particularly the visual system (Bayer, Ferrari and Erb, 2002; Jiang *et al*., 2016; La Morgia *et al*., 2016).

Visual dysfunction is increasingly recognized as an early feature of AD, preceding the onset of cognitive symptoms (London, Benhar and Schwartz, 2013; Cerquera-Jaramillo *et al*., 2018). The retina, an accessible extension of the CNS, provides a unique window to assess the brain pathology (Chow and Lang, 2001; London, Benhar and Schwartz, 2013). Structural and functional retinal abnormalities have been reported in AD patients, including thinning of the retinal nerve fiber layer (RNFL) and ganglion cell layer (GCL), reduced contrast sensitivity, and altered pupillary responses (Kim and Kang, 2019; Kremen *et al*., 2019; Risacher *et al*., 2020). Importantly, some of these retinal alterations correlate with cortical Aβ burden and cognitive impairment (on behalf of the FACEHBI study group *et al*., 2020; Song *et al*., 2021), suggesting that retinal changes may mirror early neurodegenerative processes occurring in the brain. The retina is among the most metabolically active tissues in the body and is highly dependent on mitochondrial function and efficient nutrient transport to maintain visual processing (Wong-Riley, 2010; Country, 2017; Hanna *et al*., 2022). This high bioenergetic demand may render retinal tissue particularly sensitive to early metabolic and mitochondrial disturbances that characterize AD pathogenesis (Yaffe, 2004; Dumont *et al*., 2009; Yang *et al*., 2022; Zhang *et al*., 2025).

To date, studies investigating retinal involvement in AD have largely emphasized structural and functional assessments, such as optical coherence tomography and electroretinography (Blanks *et al*., 1996; Koronyo-Hamaoui *et al*., 2011; Kim and Kang, 2019; Asanad *et al*., 2021). However, early molecular alterations in the retina during AD progression remain poorly characterized. Furthermore, it remains unclear if the retina shows similar early biochemical alterations as the brain. To this end, we investigated early proteomic changes in the retina and brain using triple-transgenic AD (3×Tg-AD) mouse model, harboring PS1^M146V^, APP^Swe^, and tau^P301L^ mutations (Oddo *et al*., 2003). In this model, detectable brain pathology typically emerges around 4-6 months of age, with intracellular Aβ accumulation beginning at 3-4 months and extracellular plaques appearing at 6-9 months, whereas 1-month-old mice do not yet show functional deficits (Belfiore *et al*., 2019; Javonillo *et al*., 2021). We therefore focused on 1-month-old 3×Tg-AD mice to capture early, pre-symptomatic alterations. We assessed retinal morphology and performed comprehensive quantitative proteomic analyses on both retina and brain tissues to identify early proteomic signatures in AD.

## Methods

### Animals

Female 3xTg-AD on a B6;129 genetic background (JAX #034830) and age-matched wild-type (WT) controls on a B6;129 background (JAX #101045) were purchased from the Jackson Laboratory. Mice were housed in the West Virginia University vivarium under a 12-hour light/12-hour dark cycle with food and water available *ad libitum*. All animal procedures were conducted in compliance with the National Institutes of Health guidelines and ARVO Statement for the Use of Animals in Ophthalmic and Vision Research. Experimental protocols were approved by the Institutional Animal Care and Use Committee (IACUC) of West Virginia University. Animals were genotyped by Transnetyx using polymerase chain reaction (PCR) analysis of genomic DNA isolated from tail samples.

### Hematoxylin and eosin staining and analysis

Mice were sacrificed at 4 weeks (postnatal day 30) and 28 weeks of age (postnatal day 200) and their eyes were carefully enucleated and immediately fixed in 1 mL of Excalibur’s Alcoholic Z-Fix (Excalibur Pathology Inc, Norman, OK). Tissues were subsequently processed, paraffin-embedded, sectioned, and stained with hematoxylin and eosin (H&E) by Excalibur Pathology. H&E-stained retinal sections were imaged using a Nikon Eclipse Ti microscope equipped with a DS-Ri2 camera (Nikon Instruments, Melville, NY). Thickness of the outer segments (OS), inner segments (IS), outer nuclear layer (ONL), and inner nuclear layer (INL) was quantified using ImageJ at six predefined locations across the retina near the optic nerve head (ONH) on both the inferior and superior sides of the eye, as previously described (Grenell *et al*., 2019).

### Tissue collection and protein extraction

Mice at 4 weeks of age were euthanized, and the eyecups were immediately enucleated (N=4 eyecups per group). Retinal dissections were performed in cold Hanks’ Balanced Salt Solution (HBSS) to isolate the neural retina, as previously described (Zhu *et al*., 2018; Xu *et al*., 2020). Whole brains were excised and briefly rinsed in cold Phosphate Buffered Saline (PBS). All tissues were flash-frozen in liquid nitrogen and stored at -80°C until further processing. For protein extraction, retina and brain samples were homogenized in radioimmunoprecipitation assay (RIPA) buffer supplemented with protease and phosphatase inhibitors (5 mg/mL). Homogenates were centrifuged at 12,000 rpm for 15 min at 4 °C, and the supernatants containing soluble proteins were collected. Protein concentrations were determined using the Pierce™ BCA Protein Assay Kit (see **Table 1**. **Key resources**).

### Quantitative proteomics

Total proteins from each sample were reduced, alkylated, and purified by chloroform/methanol extraction prior to digestion with sequencing grade modified porcine trypsin (Promega). Tryptic peptides were then separated by reverse phase XSelect CSH C18 2.5 um resin (Waters) on an in-line 150 x 0.075 mm column using an UltiMate 3000 RSLCnano system (Thermo). Peptides were eluted using a 60 min gradient from 98:2 to 65:35 buffer A:B ratio (Buffer A contains 0.1% formic acid, 0.5% acetonitrile. Buffer B contains 0.1% formic acid, 99.9% acetonitrile). Eluted peptides were ionized by electrospray (2.2 kV) followed by mass spectrometric analysis on an Orbitrap Exploris 480 mass spectrometer (Thermo). To assemble a chromatogram library, six gas-phase fractions were acquired on the Orbitrap Exploris with 4 m/z DIA spectra (4 m/z precursor isolation windows at 30,000 resolution, normalized AGC target 100%, maximum inject time 66 ms) using a staggered window pattern from narrow mass ranges using optimized window placements. Precursor spectra were acquired after each DIA duty cycle, spanning the m/z range of the gas-phase fraction (i.e. 496-602 m/z, 60,000 resolution, normalized AGC target 100%, maximum injection time 50 ms). For wide-window acquisitions, the Orbitrap Exploris was configured to acquire a precursor scan (385-1015 m/z, 60,000 resolution, normalized AGC target 100%, maximum injection time 50 ms) followed by 50x 12 m/z DIA spectra (12 m/z precursor isolation windows at 15,000 resolution, normalized AGC target 100%, maximum injection time 33 ms) using a staggered window pattern with optimized window placements. Precursor spectra were acquired after each DIA duty cycle.

### Proteomics data processing

Following data acquisition, data were searched using an empirically corrected library and a quantitative analysis was performed to obtain a comprehensive proteomic profile. Proteins were identified and quantified using EncyclopeDIA (Searle *et al*., 2018) and visualized with Scaffold DIA using 1% false discovery thresholds at both the protein and peptide level. Protein exclusive intensity values were assessed for quality using an in-house ProteiNorm (Graw *et al*., 2020), a tool for systematic evaluation of normalization methods, imputation of missing values and comparisons of multiple differential abundance methods. Normalization methods evaluated included log2 normalization (Log2), median normalization (Median), mean normalization (Mean), variance stabilizing normalization (VSN) (Huber *et al*., 2002), quantile normalization (Quantile) (Bolstad *et al*., 2003), cyclic loess normalization (Cyclic Loess) (Ritchie *et al*., 2015), global robust linear regression normalization (RLR), and global intensity normalization (Global Intensity) (Chawade, Alexandersson and Levander, 2014). The individual performance of each method was evaluated by comparing of the following metrices: total intensity, pooled intragroup coefficient of variation (PCV), pooled intragroup median absolute deviation (PMAD), pooled intragroup estimate of variance (PEV), intragroup correlation, sample correlation heatmap (Pearson), and log2-ratio distributions. The normalized data were used to perform statistical analysis using linear models for microarray data (limma) with empirical Bayes (eBayes) smoothing to the standard errors (Ritchie *et al*., 2015). Proteins with an FDR adjusted p-value < 0.05 and a fold change (FC) > 2 were considered significant. Data are available via PRoteomic IDEntifications Database (PRIDE) with identifier PXD069995.

### Bioinformatics analysis

Proteomics quantification results were further processed using bioinformatic platforms. Principal Component Analysis (PCA) and Variable Importance in Projection (VIP) scoring were performed using MetaboAnalyst (https://www.metaboanalyst.ca). Functional enrichment analysis of biological processes and cellular components for significantly changed proteins between 3xTg-AD and WT tissues, were conducted using the DAVID Bioinformatics Resources (https://davidbioinformatics.nih.gov). Protein-protein interaction network analysis was carried out using Metascape (https://metascape.org). Heatmaps and quantitative data visualizations were generated using GraphPad Prism (v9.5.1, GraphPad Software Inc). To identify AD-associated proteins enriched in the human brain proteomic dataset, we queried the Agora AD Knowledge Portal (https://agora.adknowledgeportal.org). Identified associations were further validated through manual curation of published literature using PubMed, with combinations of relevant keywords (e.g., “[protein name] AND [Alzheimer’s disease OR neurodegenerative disease]”).

### Statistics

The statistical analyses of retinal thickness are presented as the mean ± standard deviation (SD). Unpaired two-tailed t tests were performed in GraphPad Prism v9.4.1 to determine significance. Values p < 0.05 were considered significant.

## Results

### Age-dependent retinal thinning in 3xTg-AD mice

To examine retinal morphological changes, we quantified the thickness of individual retinal layers in H&E-stained sections at postnatal day 30 (P30) and postnatal day 200 (P200). At P30, retinal structure in 3xTg-AD mice was comparable to that of WT controls, without differences across retinal layers (**Figure 1A, 2A, C–F**). However, multiple retinal layers including OS, IS, ONL, and INL were significantly shorter in 3xTg-AD mice at P200 (**Figure 2G–J**). This progressive decline in retinal thickness with age in 3xTg-AD mice is consistent with reports of retinal layer thinning in AD patients (Pillai *et al*., 2016; Cunha *et al*., 2017; Kim and Kang, 2019).

**Figure 1.**
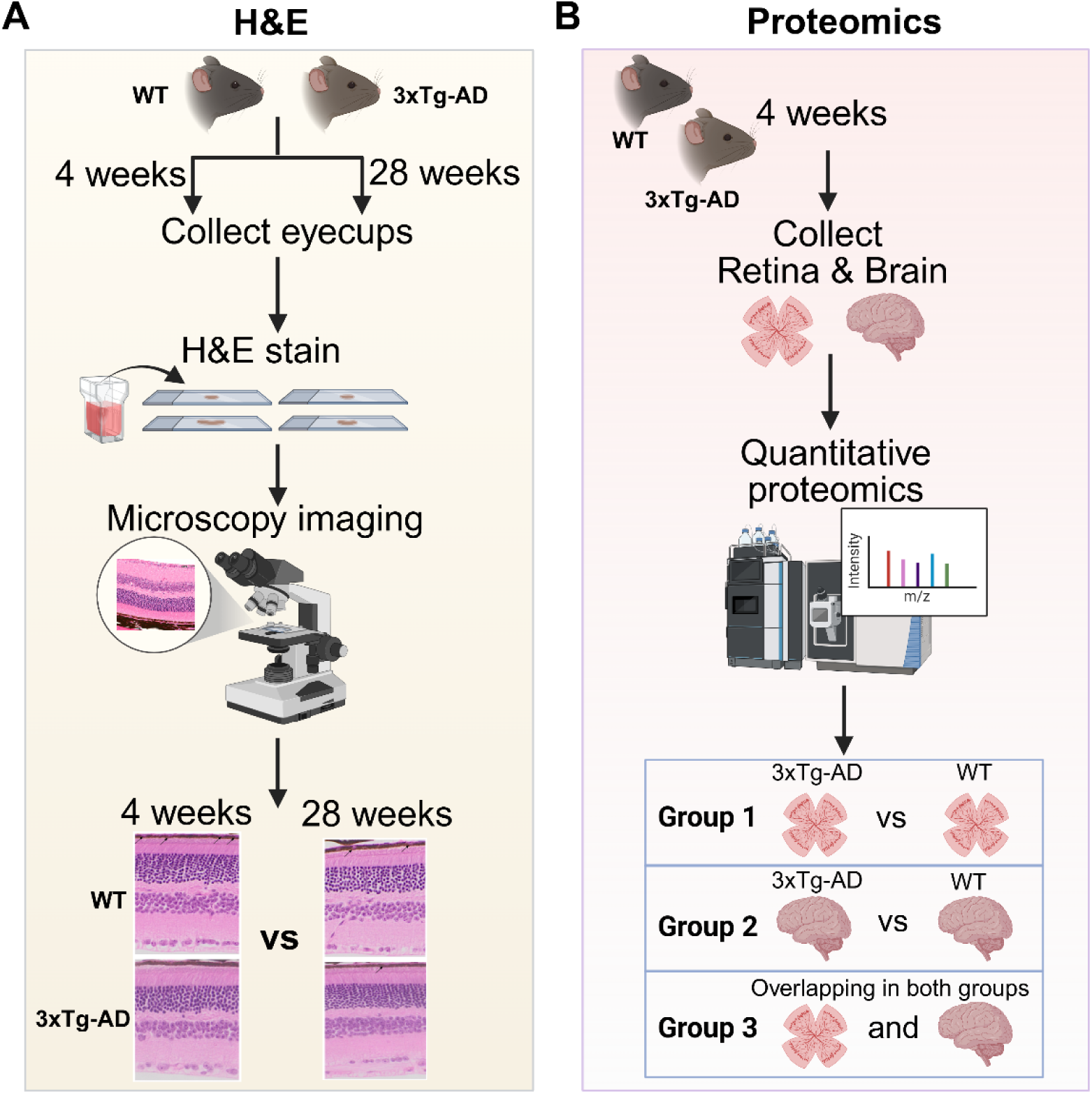
Experimental workflow. **(A)** Retinal morphology was assessed by Hematoxylin and Eosin (H&E) staining of eyecup sections from 3xTg-AD mice and WT controls at 4- and 28-weeks of age. **(B)** Quantitative proteomics was performed on retina and brain tissues from 4-weeks old 3xTg-AD mice and WT controls.

**Figure 2.**
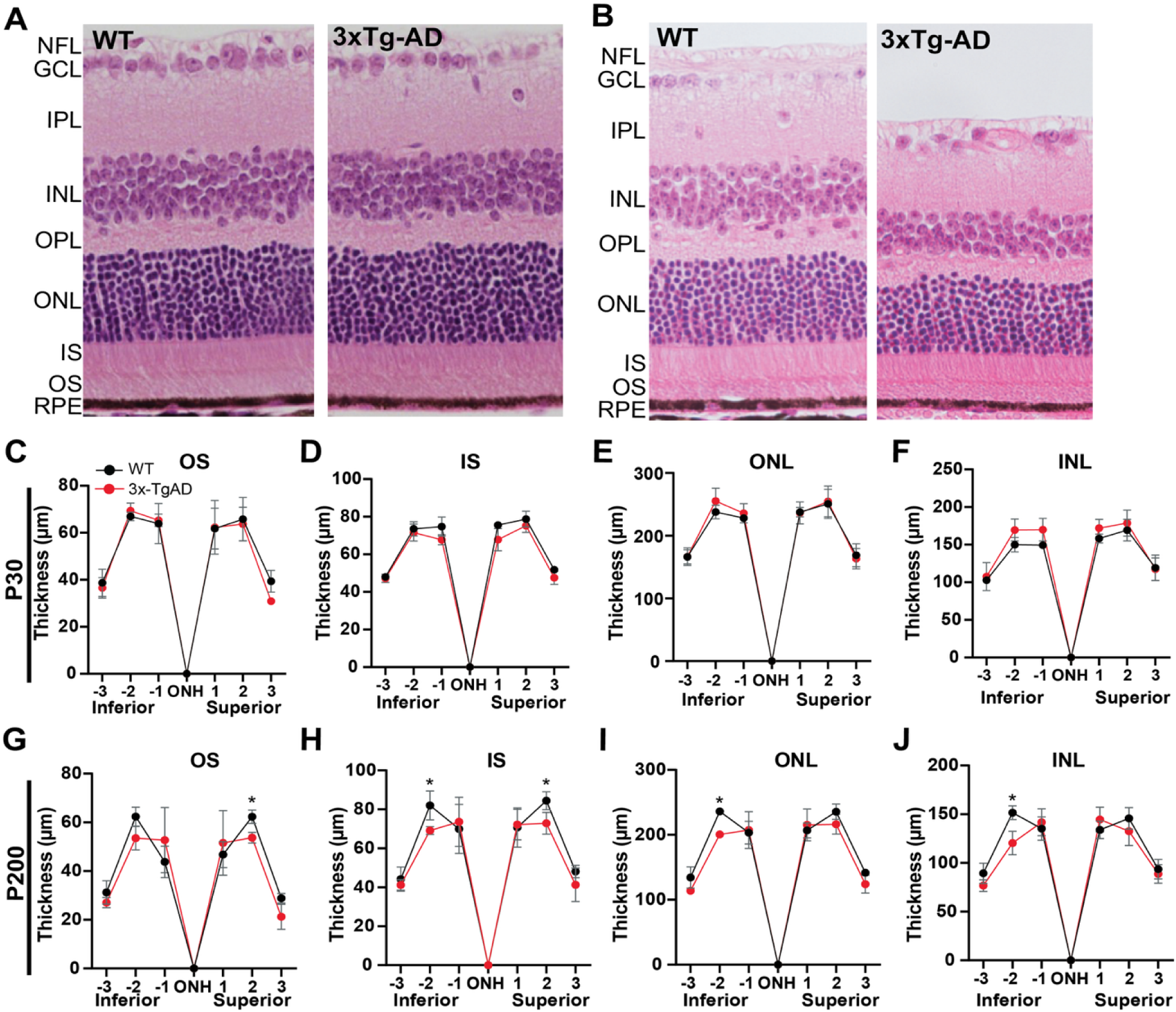
Age-associated structural alterations in the retina of 3xTg-AD mice. Representative H&E-stained retinal sections from WT and 3xTg-AD mice at **(A)** P30 and **(B)** P200. **(C–F)** Quantification of retinal layer thickness at P30 at the indicated positions from the optic nerve head (ONH). **(G–J)** Quantitative assessment of the thickness of retinal layers thickness at P200. Asterisks indicate significant differences determined through *t* tests between the WT and 3xTg-AD groups. Data is presented as mean ± SD. RPE, retinal pigment epithelium; OS, outer segment; IS, inner segment; ONL, outer nuclear layer; INL, inner nuclear layer; IPL, inner plexiform layer; GCL, ganglion cell layer; NFL, Nerve fiber layer. N = 4 eyecups/group.

### Early retinal proteomic alterations in 3xTg-AD mice

To investigate early proteomic alterations preceding AD pathology, we performed quantitative proteomic analysis of retinas from 3xTg-AD and WT mice at P30 (**Figure 1B**). Principal component analysis (PCA) showed a clear separation between two groups, indicating early global proteomic change in the retina from 3xTg-AD mice (**Figure 3A**). A total of 92 proteins were significantly altered in 3xTg-AD retinas, including 51 downregulated and 41 upregulated proteins (**Figure 3B**; **Table S1**). Variable Importance in Projection (VIP) analysis identified top 15 proteins driving PCA group separation (**Figure 3C**) with prominent changes of mitochondrial proteins involved in energy metabolism, such as succinate-CoA ligase subunit β (SUCB2) and ADP/ATP translocase 4 (ADT4), as well as hemoglobin-related proteins (HBB1, HBB2, and A8DUK4).

**Figure 3.**
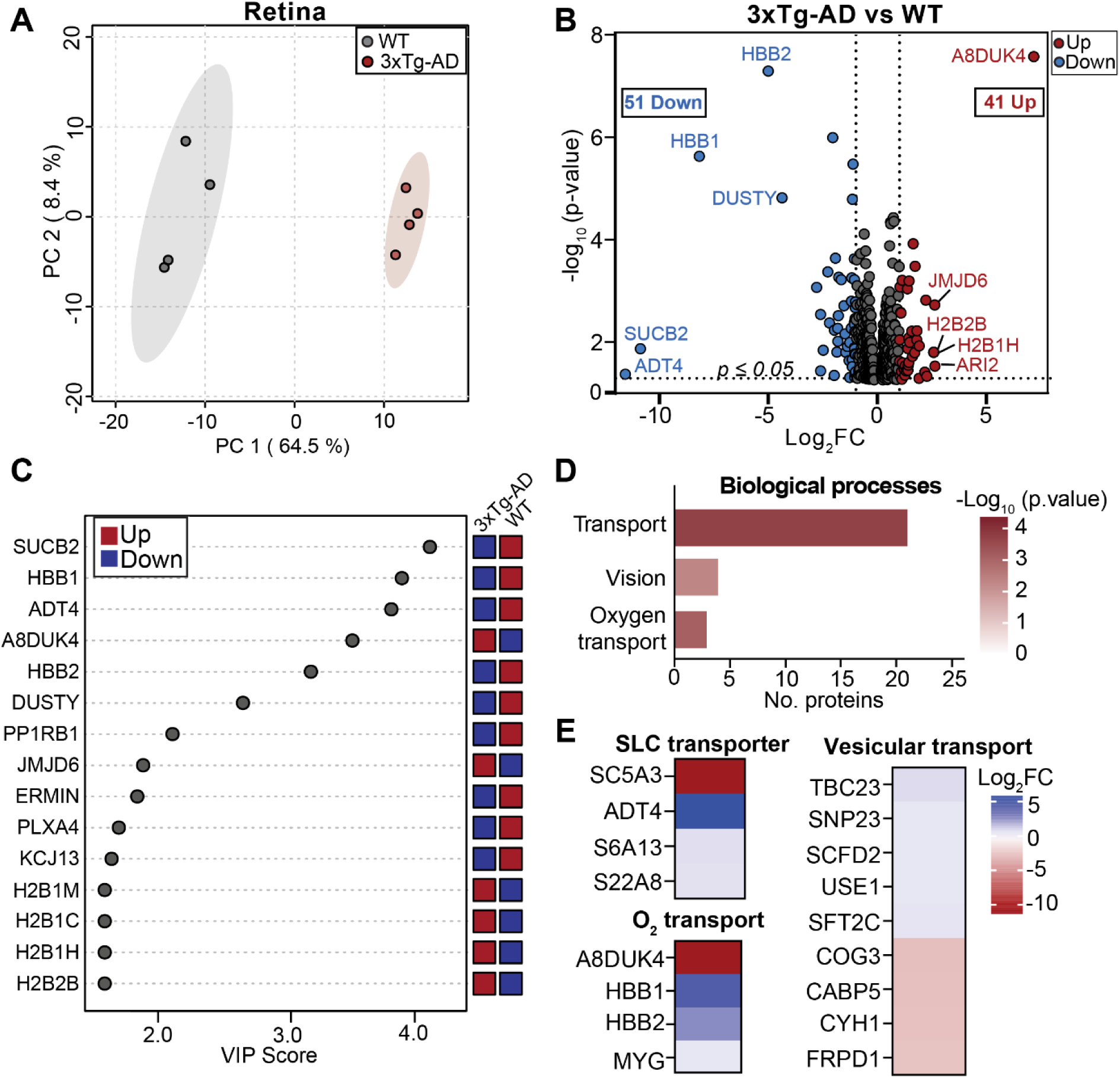
Early retinal proteomic changes in 3xTg-AD mice. **(A)** Two-dimensional principal component analysis (PCA) plot shows clear separation between 3xTg-AD and WT retina proteome. **(B)** Volcano plot of differentially expressed proteins in the retina of 3xTg-AD compared to WT. Top five up-and downregulated are labeled (|log_2_FC| ≥ 1, p ≤ 0.05; red = upregulated, blue = downregulated in 3xTg-AD). **(C)** Variable Importance in Projection (VIP) plot highlighting the top 15 proteins that separate 3xTg-AD from WT retina. Tiles on the right indicate up-(red) and down-regulated (blue). **(D)** Biological pathways enriched in 3xTg-AD identified by Gene ontology (GO) analysis. **(E)** Heatmaps of transport-related processes with representative proteins enriched in 3xTG-AD versus WT. SLC, solute carrier transporter. N=4.

Specifically, proteins associated with retinal structure and neuronal connectivity, including ermin (ERMIN) and plexin-A4 (PLXA4), were decreased, suggesting early neuronal vulnerability. Multiple histone H2B variants (H2B1M, H2B1C, H2B1H, H2B2B) and the histone demethylase JMJD6, however, were increased (**Figure 3C**), indicating early epigenetic alterations and chromatin remodeling. Gene ontology (GO) analysis showed enrichment of pathways related to transport, in particular oxygen transport, and vision (**Figure 3D, E; Table S2**). Several solute carrier (SLC) transporters were differentially expressed in 3xTg-AD retina. SC5A3 was significantly upregulated, whereas ADT4, S6A13, and S22A8 were downregulated (**Figure 3E**). Notably, ADT4, a mitochondrial ADP/ATP translocase, was reduced by more than 11-fold, suggesting impaired mitochondrial energy output. In addition, proteins involved in vesicular transport were broadly altered. Components associated with endoplasmic reticulum (ER)-Golgi trafficking and synaptic vesicle dynamics, including TBC23, SNP23, SCFD2, USE1, and SFT2C, were decreased. In contrast, expression of proteins involved in Golgi organization, calcium-dependent synaptic signaling, and vesicle recycling such as COG3, CABP5, CYH1, and FRPD1, was increased. Strikingly, vision-related proteins essential for photoreceptor maintenance and retinal signaling, including RGR, RDH5, RPE65, IMPG2, and KCNJ13, were uniformly downregulated (**Figure S1**). Together, these findings demonstrate early retinal proteomic changes in 3xTg-AD mice, characterized by chromatin remodeling, disrupted energy metabolism, intracellular transport, and vision-associated pathways.

### Coupled mitochondrial and nuclear protein alterations in the AD retina

Building on the pathway-level dysregulation identified in early AD retinas, we next examined the subcellular localization of differentially expressed proteins. Cellular component enrichment analysis demonstrated that significantly altered proteins in 3xTg-AD retinas were predominantly localized to the nucleosome core, mitochondria, and chromosome (**Figure 4A; Table S2**). Multiple mitochondrial proteins involved in metabolism and energy production, including LACB2, LPIN1, SRAC1, NDUB2, ARGI1, SUCB2, and ADT4, were downregulated in the AD retina (**Figure 4B**). In contrast, proteins associated with nuclear processes were upregulated, including histone variants and factors involved in DNA replication (RFC5), RNA processing (RNH2A, DDX55), and chromatin remodeling (SMRCD).

**Figure 4.**
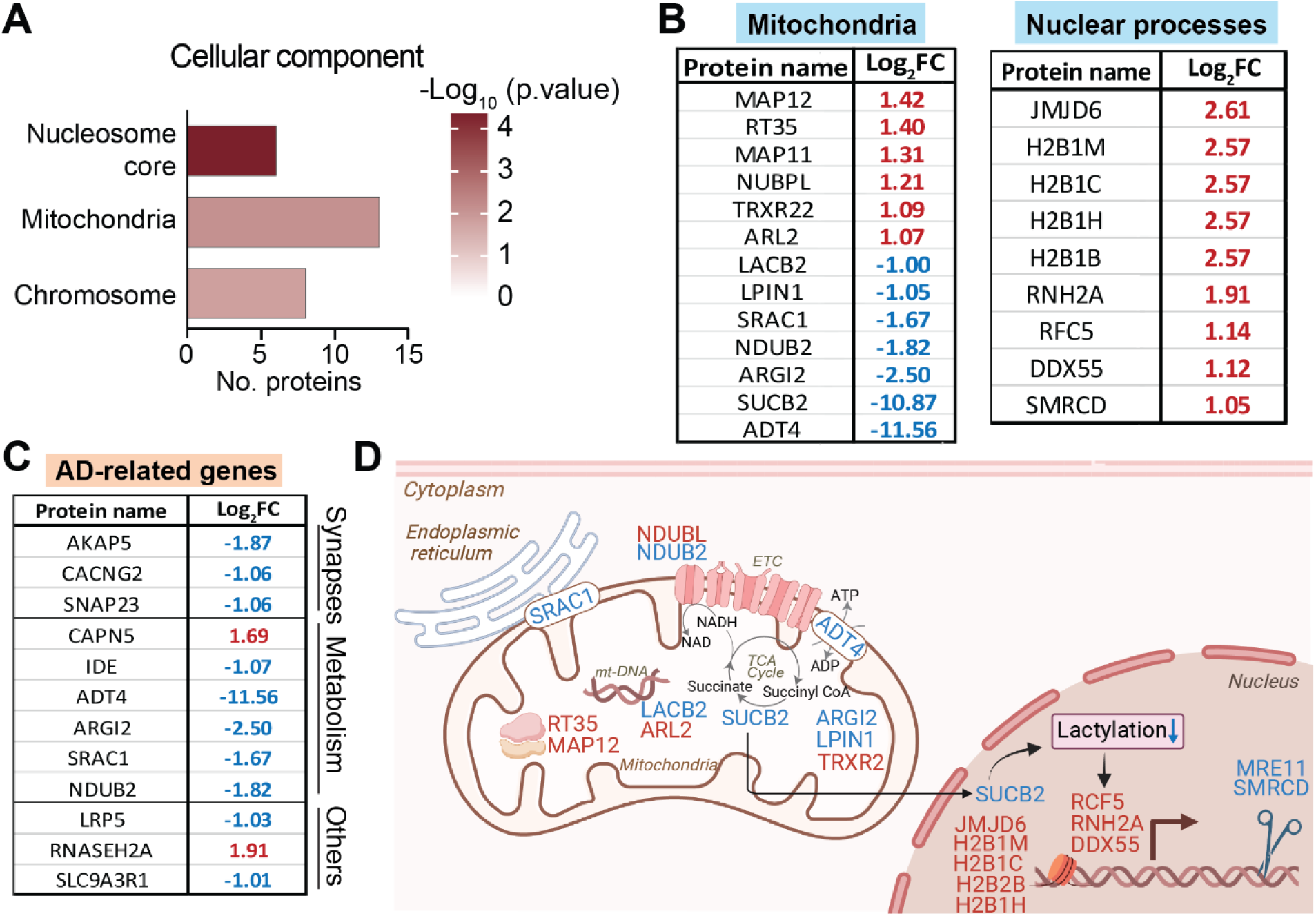
Differential mitochondrial and nuclear proteome changes in 3xTg-AD retina. **(A)** GO enrichment for cellular components altered in 3xTg-AD retina. **(B)** Differentially expressed mitochondria-and nucleus-related proteins in the retina (red = upregulated, blue = downregulated in 3xTg-AD vs WT). **(C)** Proteins differentially expressed in 3xTg-AD retina that are linked to AD pathology, identified using Agora AD Knowledge Portal and manual literature search. **(D)** Proposed working model illustrating how mitochondrial proteome changes may signal to the nucleus and reprogram nuclear gene expression in the retina. See **Table S1** for detailed protein annotations and fold changes. AD, Alzheimer’s disease; mt-DNA, mitochondrial DNA; ETC, electron transport chain; TCA cycle, tricarboxylic acid cycle.

To determine whether the significantly altered retinal proteins identified in 3xTg-AD mice have known associations with AD in humans, we queried the Agora AD Knowledge Portal to identify proteins linked to AD pathogenesis and to explore potentially shared pathways. We found that twelve of significantly altered retinal proteins involved in synaptic function and metabolism have prior evidence linked to AD pathogenesis (**Figure 4C**). In particular, several mitochondrial proteins which are changed in the 3xTG-AD retina (e.g. ADT4, SRAC1, ARGI2), were also altered in AD patients. Collectively, the reduction of mitochondrial proteins together with increased chromatin-associated factors supports a model in which early retinal stress in AD is driven by mitochondrial dysfunction (**Figure 4D**). In particular, reduced expression of SUCB2, a key enzyme involved in protein succinylation and lactylation, may further perturb mitochondria-to-nucleus communication by modifying chromatin-associated posttranslational marks and thereby modulating metabolic gene expression (Hu *et al*., 2023; Liu *et al*., 2025; Tsusaka *et al*., 2025). Together, these findings indicate that mitochondrial dysfunction and associated nuclear responses represent an early molecular feature of retinal involvement in AD.

### Early proteomic alterations in 3xTg-AD brain

Multivariate PCA demonstrated a clear separation between 3xTg-AD and WT brain proteomes, indicating substantial global differences in protein expression (**Figure 5A**). Differential expression analysis identified 130 significantly altered proteins, including 38 downregulated and 92 upregulated in the AD brain (**Figure 5B**; **Table S3**). Similar to the retina, hemoglobin subunits (HBB1, HBB2, and A8DUK4) were among the most prominently altered proteins. GO analysis of biological processes showed significant enrichment of pathways related to molecular transport and the mitochondrial respiratory chain in the 3xTg-AD brain (**Figure 5C; Table S4**). Consistent with this, cellular component analysis indicated that differentially expressed proteins were primarily localized to mitochondria, membrane, cytoplasmic vesicles, and synaptic compartments (**Figure 5D; Table S4**). Assessment of mitochondrial pathways showed broad changes in proteins involved in oxidative phosphorylation and electron transport chains, with strong enrichment of subunits from Complex I and Complex III (**Figure 5E**). These included multiple NADH dehydrogenase subunits, such as NDUS1, NDUS5, NDUA6, as well as cytochrome c oxidase–associated proteins, including COX5A, CX6A1, and CX7A2.

**Figure 5.**
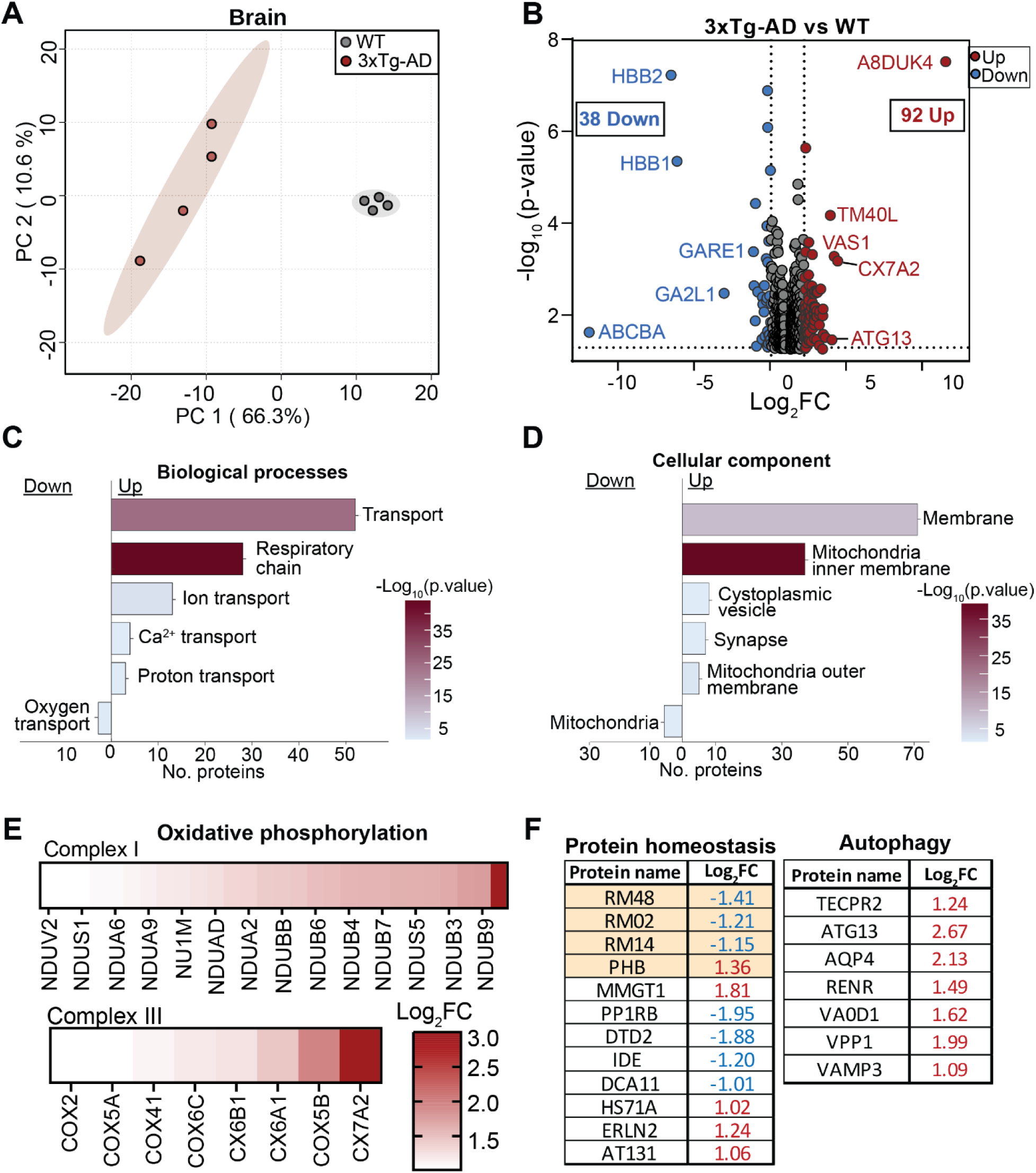
Early proteomic changes in the 3xTG-AD mouse brain. **(A)** Two-dimensional PCA plot shows clear separation between 3xTg-AD and WT brain proteome. **(B)** Volcano plot of differentially expressed proteins in the brain of 3xTg-AD compared to WT. Top five up- and downregulated are shown (|log2FC| ≥ 1, p ≤ 0.05; red = upregulated, blue = downregulated in 3xTg-AD). **(C)** GO enrichment analysis for biological processes and **(D)** cellular components altered in 3xTg-AD brain. **(E)** Enrichment of oxidative phosphorylation-related proteins in 3xTg-AD brain. **(F)** Differentially expressed 3xTg-AD brain proteins linked to protein homeostasis and autophagy, identified by PubMed literature search. Mitochondrial proteins are highlighted in orange. N=4.

In addition to respiratory chain alterations, proteins involved in mitochondrial protein homeostasis and autophagy were also differentially expressed (**Figure 5F**). These included downregulation of ribosomal and mitochondrial-associated proteins such as large ribosomal subunit protein mL48 (RM48), uL2m (RM02), uL14m (RM14), and prohibitin 1 (PHB), as well as upregulation of autophagy-related proteins including ATG13 and TECPR2. Protein–protein interaction network analysis further highlighted clustering of altered proteins within pathways related to oxidative phosphorylation, mitophagy, and synaptic vesicle maturation (**Figure S2**). Taken together, these findings indicate that proteomic alterations in the 3xTg-AD brain are dominated by changes in mitochondrial-associated proteins, particularly those involved in respiratory chain function and mitochondrial protein homeostasis.

### Comparison of early proteomic alterations in retina and brain in 3xTg-AD mice

To investigate how early proteomic changes were shared between the retina and brain, we compared proteins significantly altered in each tissue of 3xTg-AD mice relative to WT controls. Among the 92 proteins altered in the retina and the 130 in the brain, eight proteins were significantly changed in both tissues (**Figure 6A**). These shared proteins included three hemoglobin-related proteins (HBB1, HBB2, and A8DUK4), and several proteins involved in metabolic regulation and protein turnover, including insulin-degrading enzyme (IDE), lactoylglutathione lyase (LGUL), E3 ubiquitin-protein ligase (PP1RB), TBC1 domain family member 23 (TBC23), and zinc finger CCCH-type containing protein 7B (F8VPP8). Except for A8DUK4 and F8VPP8, all overlapping proteins were reduced in both tissues (**Figure 6B**).

**Figure 6.**
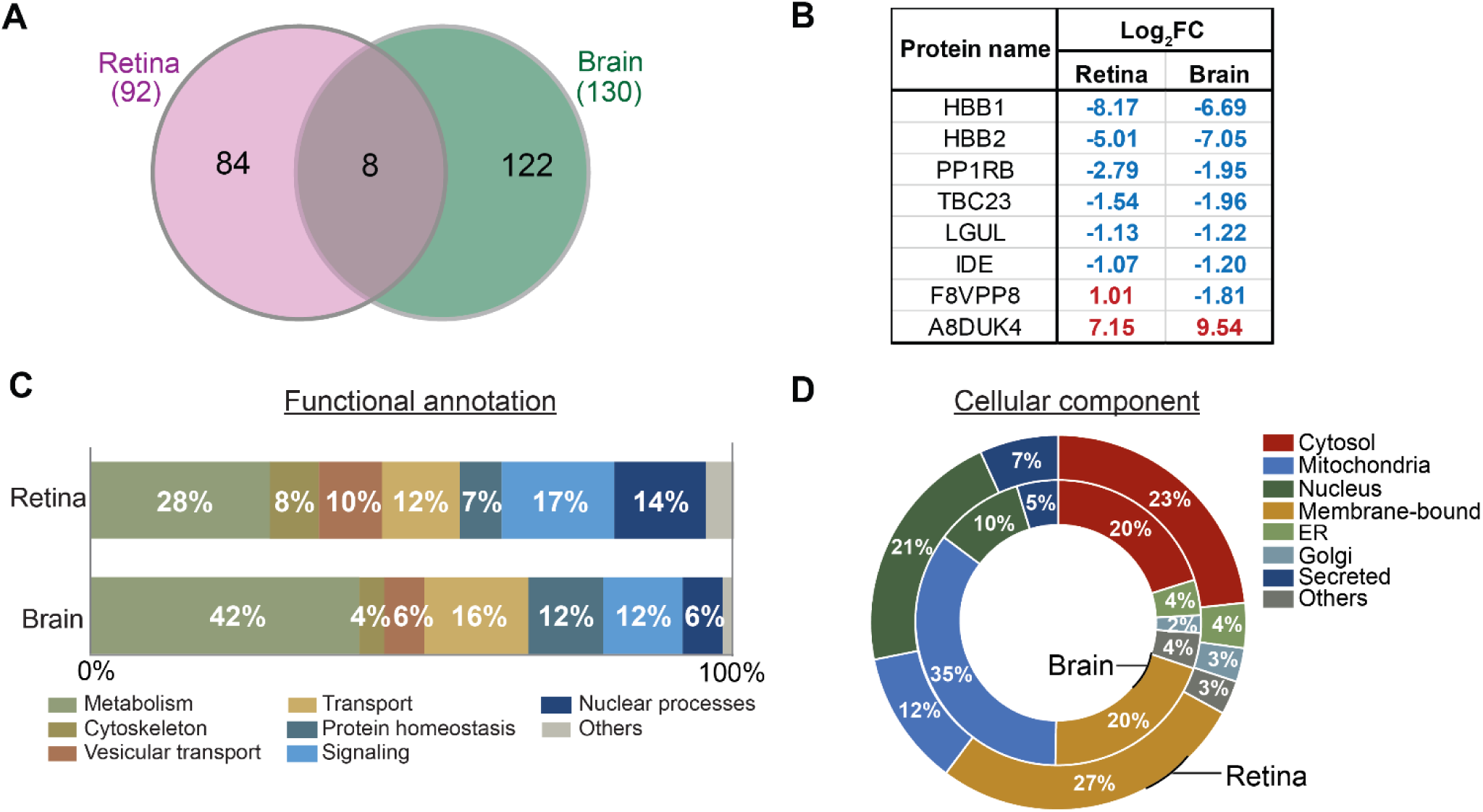
Comparison of differentially expressed proteins in retina and brain. **(A)** Total significantly changed and overlapping proteins in retina and brain. **(B)** Overlapping proteins between retina and brain. **(C)** Bar plots represent the percentage distribution of significantly changed proteins based on functional annotation and **(D)** cellular components in retina (outer plot) and brain (inner plot). ER, Endoplasmic Reticulum; Membrane-bound, includes proteins in cellular and intracellular membranes.

Pathway and cellular localization analyses further revealed common changes across the two tissues. GO analysis showed that metabolism, transport, and signaling were among the most highly enriched biological processes in both retina and brain (**Figure 6C**). Consistently, differentially expressed proteins were predominantly localized to cytosolic, membrane-associated, mitochondrial, and nuclear compartments (**Figure 6D**). Notably, the AD brain showed a greater proportion of changes in mitochondrial proteins, whereas membrane-associated proteins were more enriched in the retina. Together, these comparative analyses indicate that the retina and brain share early AD-associated proteomic alterations while also have tissue-specific changes during initial stages of disease.

## Discussion

In this study, we have identified early proteomic signatures in the retina and brain of 3xTg-AD mice. Although only eight proteins overlapped between the two tissues, both showed similar alterations in mitochondrial metabolism, oxygen, and intracellular transport. Besides common changes, the retinal proteome showed specific alterations in vision-related, transporters, and chromatin-associated proteins, whereas the brain proteome featured changes in mitochondrial respiratory chain components and protein homeostasis. Many of these dysregulated pathways are recognized as progressive drivers or early features of AD in patients (Decker *et al*., 2010; Di Filippo *et al*., 2010; Kundra *et al*., 2017; Kazemeini *et al*., 2024) (Koronyo *et al*., 2023). Our findings further support the retina as a sensitive “window to the brain” for detecting preclinical AD pathology.

Among the shared alterations, β-globin subunits HBB1, HBB2, and A8DUK4 stand out as consistently dysregulated in both tissues. While hemoglobin is classically associated with erythrocytes, neurons and glial cells also express hemoglobin to facilitate oxygen uptake under hypoxic stress (Biagioli *et al*., 2009; Schelshorn *et al*., 2009). Altered hemoglobin expression and localization have been reported in AD brain, including enrichment near Aβ plaques and cerebral amyloid angiopathy (Wu *et al*., 2004; Ashraf, Dani and So, 2020), as well as paradoxical depletion in neurons containing neurofibrillary tangles (Ferrer *et al*., 2011; Altinoz *et al*., 2019). Beyond oxygen transport, neuronal hemoglobin has been implicated in mitochondrial function, mitochondria-to-nucleus signaling, epigenetic regulation, and autophagy (Ferrer *et al*., 2011; Codrich *et al*., 2017; Altinoz *et al*., 2019). In addition, hemoglobin can localize to the inner mitochondrial membrane (Shephard *et al*., 2014), interact with ATP synthase (Brown *et al*., 2016), and, enhance mitochondrial respiration and improves neuronal oxygen utilization in dopaminergic neurons (Biagioli *et al*., 2009; Schelshorn *et al*., 2009). In contrast to the brain, the role of hemoglobin in the neural retina remains unclear. Human retinal pigment epithelial (RPE) cells can synthesize and secrete hemoglobin (Tezel *et al*., 2009). In addition, hemoglobin expression in retinal macroglia and ganglion cells increases under hypoxic conditions to support cell survival (Tezel *et al*., 2010). Given the high metabolic demand and relatively low physiological oxygen availability in the retina (Hurley, 2021), locally expressed hemoglobin may help preserve the mitochondrial function. Altered expression of hemoglobin subunits in both retina and brain at this early stage of AD may therefore reflect a compensatory adaptation to mitochondrial dysfunction and oxidative stress. However, how neuronal hemoglobins contribute to AD pathogenesis requires further investigation.

Mitochondrial dysfunction is a key contributor in AD pathogenesis, with extensive evidence of impaired oxidative metabolism, reduced ATP production, altered tricarboxylic acid (TCA) cycle, and respiratory chain activity (Onyango, Dennis and Khan, 2016; Buchman *et al*., 2019; Wang *et al*., 2020, 2024). Key metabolic enzymes SUCB2 and ADT4 were reduced more than 10- and 11-folds, respectively, in the 3xTg-AD retina (**Figure 4B**). SUCB2, encoded by *SUCLG2*, catalyzes the reversible conversion of succinyl-CoA to succinate coupled with GTP synthesis (Phillips *et al*., 2009). This reaction is particularly important in the retina, where GTP is required for cGMP production during phototransduction (Du *et al*., 2016). Beyond its metabolic role, SUCB2 is important for protein succinylation and lactylation, linking metabolism to epigenetic regulation (**Figure 4D**) (Zhang *et al*., 2023; Liu *et al*., 2025). Altered succinylation of mitochondrial proteins, APP, and tau has been reported in human AD brains and mouse models (Pan *et al*., 2022; Yang *et al*., 2022; Zhang *et al*., 2025). Similarly, ADT4 (*SLC25A31*), a mitochondrial ADP/ATP translocase, responsible for exchanging matrix ATP with cytosolic ADP across the inner mitochondrial membrane, is massively reduced. Such impaired ADP/ATP exchange can compromise oxidative phosphorylation, alter mitochondrial membrane potential, and increase susceptibility to cellular stress (Clémençon, Babot and Trézéguet, 2013; Gutiérrez-Aguilar and Baines, 2013; Jia and Du, 2021). The substantial reduction of SUCB2 and ADT4 in early AD retina would limit the availability of GTP and ATP, compromising mitochondrial metabolism and potentially affecting phototransduction and metabolite-driven signaling. It warrants future investigation whether these metabolic defects contribute to downregulation of retinal proteins involved in the visual cycle, ion transport, and photoreceptor maintenance before visual impairment in 3xTg-AD mice and AD patients (Kusne *et al*., 2017; Kim and Kang, 2019). These findings further support that mitochondrial dysfunction emerges early in the retina in AD pathogenesis.

This study has several limitations. We examined a single mouse model at one early time point, which provides a snapshot of early proteomic changes but does not capture the progression of these changes as pathology advances. Longitudinal studies are needed to track these proteomic alterations over the course of disease. While we identified candidate pathways based on altered proteins, functional studies are required to establish causal relationships between specific protein changes and tissue dysfunction. Furthermore, the 3xTg-AD model does not fully represent the complex genetic, environmental, and sporadic factors that characterize human AD. Validation in human retinal and brain tissues will therefore be essential to assess the translational relevance of these findings. Despite these limitations, our study provides the first comprehensive proteomic characterization of both retina and brain in the 3xTg-AD model, demonstrating early metabolic, mitochondrial, chromatin-related, and transport alterations that precede detectable pathology. These findings establish a molecular foundation for understanding early AD mechanisms and support the identification of retinal-based biomarkers for early detection of disease.

## Author Contributions

Conceptualization: Jianhai Du

Methodology: Artjola Puja, Rachel McNeel

Investigation (experiments/data collection): Artjola Puja, Rachel McNeel, Rong Xu

Resources (animals/samples/instrumentation): Artjola Puja, Rachel McNeel, Rong Xu

Visualization (figures/tables): Artjola Puja, Rachel McNeel

Writing – Review & Editing: Artjola Puja, Rachel McNeel, Jianhai Du, David Hansman

Supervision: Jianhai Du

Funding Acquisition: NIH Grants (EY026030, EY031324, EY032462), the Retina Research Foundation, NIH/NIGMS grant R24GM137786 to IDeA National Resource for Quantitative Proteomics, NIH NIGMS P20GM144230 Visual Sciences COBRE grant to WVU, and an unrestricted challenge grant from Research to Prevent Blindness (RPB) to the Ophthalmology department at WVU.

## Acknowledgements

This work was supported by NIH Grants (EY026030, EY031324, EY032462), the Retina Research Foundation, NIH/NIGMS grant R24GM137786 to IDeA National Resource for Quantitative Proteomics, NIH NIGMS P20GM144230 Visual Sciences COBRE grant to WVU, and an unrestricted challenge grant from Research to Prevent Blindness (RPB) to the Ophthalmology department at WVU.

## Data availability

The mass spectrometry proteomics data have been deposited to the PRoteomic IDEntifications Database (PRIDE) with the dataset identifier PXD069995.

## Abbreviations

AD: (Alzheimer’s disease)
WT: (wild-type)
3x-Tg-AD: (triple-transgenic-AD)
SUCB2: (succinate-CoA ligase subunit β)
HBB: (hemoglobin subunit β)
ER: (endoplasmic reticulum)
Aβ: (amyloid-β)
GO: (Gene Ontology).

## Notes

### Competing Interest Statement

The authors have declared no competing interest.

