## Supplementary Figures for "Early proteomic signatures of Alzheimer’s disease in the retina and brain of 3xTg-AD mice"

This PDF file includes:

Supplementary Figures

Supplementary Table

**Supplementary Figures**

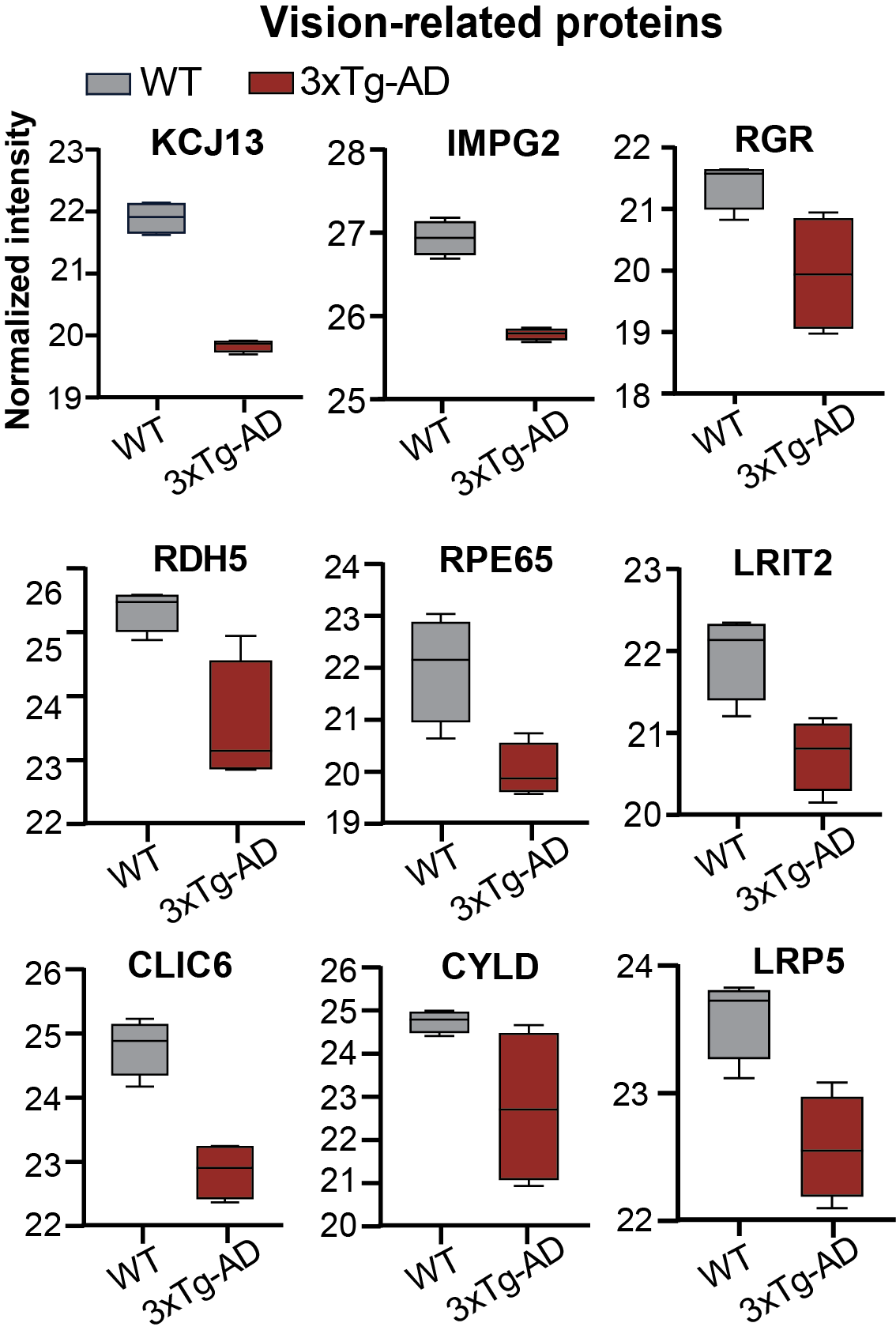

**Figure S1.** Bar charts of significantly changed vision-related proteins in the retina of 3xTg-AD compared to WT controls. N=4

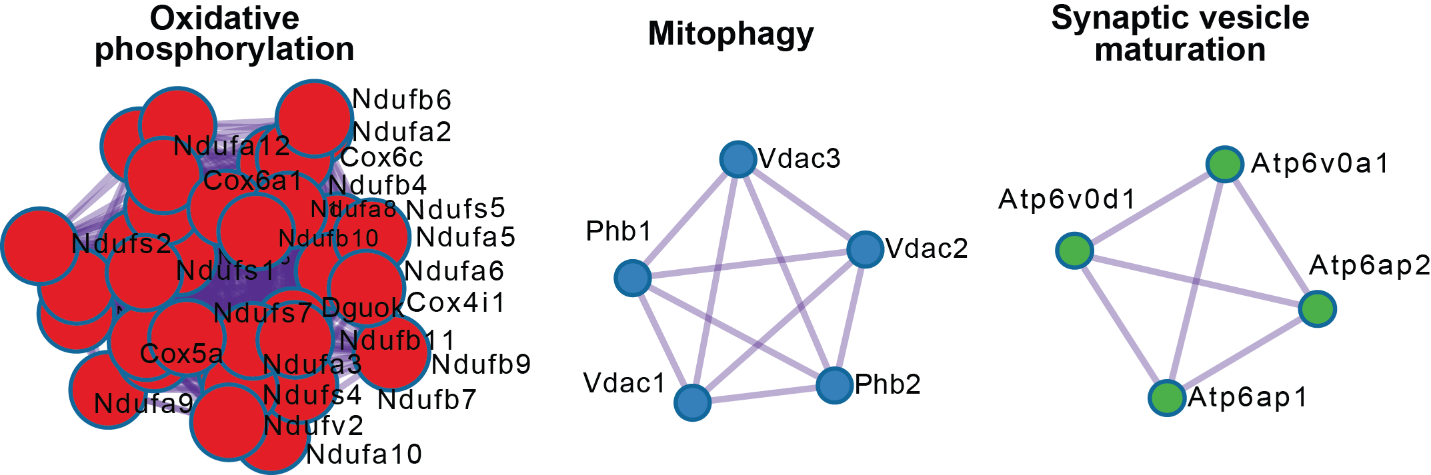

**Figure S2.** Clustering of the protein–protein interaction network of differentially expressed proteins in the brain tissues of 3xTg-AD mice. Analysis was performed using Metascape. N=4

**Table S5. Key resources**

| **Mouse Strain** | **Catalog** | **Company** | **Location** |
| --- | --- | --- | --- |
| 3xTg-AD | MMRRC_034830-JAX | The Jackson Laboratory | Bar Harbor, ME, USA |
| B6129SF1/J | IMSR_JAX:101043 | The Jackson Laboratory | Bar Harbor, ME, USA |
| **Reagents** | **Catalog** | **Company** | **Location** |
| RIPA Lysis and Extraction Buffer | 89901 | Thermo Fisher Scientific | Rockford, IL USA |
| Pierce Protease and Phosphatase Inhibitor Mini Tablets | A32959 | Thermo Fisher Scientific | Rockford, IL USA |
| Pierce™ BCA Protein Assay Kit | 23225 | Thermo Fisher Scientific | Rockford, IL USA |
| Excalibur’s Alcoholic Z-Fix | 1024 | Excalibur Pathology Inc. | Norman, OK USA |
| Phosphate Buffer Saline | 3013794 | Thermo Fisher Scientific | St. Louis, MO USA |
| Hanks’ Balanced Salt Solution | 2276814 | Thermo Fisher Scientific | Waltham, MA USA |
